# Infant gut microbiota in multi-child families converges with adults without a sibling-specific signature

**DOI:** 10.1101/2025.10.31.685886

**Authors:** Eveliina Hanski, Alise J Ponsero, Kaija-Leena Kolho, Willem M de Vos, Anne Salonen

**Affiliations:** Human Microbiome Research Program, University of Helsinki, Helsinki, Finland; University of Oxford, Oxford, UK; Collegium Helveticum, Zurich, Switzerland; Human Microbiome Research Program, Faculty of Medicine, University of Helsinki, Helsinki, Finland; Core bioinformatics, Quadram Institute of Biosciences, Norwich UK; Children’s Hospital HUS, University of Helsinki and Faculty of Medicine, University of Helsinki, Helsinki, Finland; Laboratory of Microbiology, Wageningen University, The Netherlands

## Abstract

The human gut microbiota undergoes rapid development during early life. Sibling presence has been associated with altered infant microbiota composition and accelerated maturation, but distinguishing direct microbial transmission between siblings from confounding by maternal reproductive history (parity) remains challenging. We analysed >5,000 faecal samples from >800 mother-father-infant triads in the Finnish HELMi birth cohort to characterise microbiota patterns that could inform underlying mechanisms. While sibling presence was associated with accelerated microbiota maturation and modest compositional differences, infants who both had siblings (or both lacked siblings) were no more similar to each other than mixed pairs. Instead, infants with siblings exhibited non-specific convergence with adult microbiota profiles and contributed fewer unique genera to their family metacommunities (infant–mother–father triads). Multi-child families harboured smaller but compositionally distinct metacommunities. These patterns challenge simple models of direct sibling transmission and suggest that multiple factors associated with family structure may jointly influence infant microbiota development, though specific mechanisms remain unclear.

## Introduction

The human gut microbiota is a complex and dynamic community of microorganisms that plays a crucial role in health through its involvement in nutrient absorption, immune system functioning, and metabolic regulation^1^. The early-life microbiota is particularly important because it actively contributes to the development of critical host systems^2^, such that alterations during this period can have lasting health effects that persist even after initial microbiota differences have been resolved^3^. As such, understanding how multiple factors influence microbiota development in early life is critical for both immediate and long-term health outcomes.

The establishment of the gut microbiota begins at birth and continues through early life, driven by microbial transmission as well as selective forces posed by environmental factors (e.g., delivery mode, diet, antibiotics^4^) and host physiology (e.g., immune system^5^). While initial colonisation occurs through maternal transmission from vaginal and gut microbiota^6^, ongoing microbial exchange between individuals becomes increasingly important as infants develop and interact more broadly with their environment. This social transmission process is facilitated by the highly individualised nature of the human gut microbiota: each person hosts a unique microbial community that can both influence and be influenced by the microbiota of others^7–9^. Growing evidence demonstrates that such social transmission can significantly shape microbial community development^6,10,11^.

Within this framework of social transmission, sibling presence represents a particularly important influence on infant gut microbiota development^4,12–15^. Studies have documented that sibling presence is associated with accelerated microbiota maturation^4,12^. This might be because siblings are still undergoing microbiota development themselves, creating a temporally shifting pool of microbes, which may expose the index infant to microbes from later stages of microbiota development and as such accelerate the process of microbiota maturation. Additional factors may also explain ‘sibling effects’ on infant microbiota development. For example, siblings who attend day care or school expose the index infant to microbes from a wider circle of children beyond just their immediate household^11^. Further, the social dynamics of infant–infant microbial transmission may differ from infant–adult interactions: children may share toys and engage in close physical play, creating opportunities for microbial exchange that may be distinct from interactions with adults. Finally, siblings may also influence the index child’s behaviour in ways that could affect the gut microbiota. For instance, the infant may imitate their older siblings’ behaviours, such as trying new foods, which could shape the infant’s diet^16^ and subsequently influence their gut microbiota.

However, in many populations, the presence of siblings is closely correlated with parity (number of previous late-stage pregnancies), creating a fundamental challenge in interpreting these findings. Parity and broader aspects of maternal reproductive history (including miscarriages and type of previous childbirths) may independently influence infant microbiota through distinct biological pathways. Previous pregnancies and deliveries could alter maternal immune function and hormonal profiles, potentially changing maternal microbiota composition that is then transmitted to the infant. Supporting this, parity has been shown to significantly alter both maternal vaginal and gut microbiota composition^17–19^, suggesting previous pregnancies may induce ‘ecological memory’ that modulates how the maternal microbiota responds to subsequent pregnancies. Beyond these biological pathways, parenting behaviours may differ between first- and subsequent-born children in ways that shape microbial exposures, such as more cautious hygiene practices among first-time parents^20^.

While studies have identified both maternal parity and sibling presence as potential factors influencing infant gut microbiota development, the extent to which these factors operate independently or interactively remains unclear. This distinction is particularly important because sibling-associated microbiota changes have been linked to health outcomes, namely reduced risk of allergic diseases. However, findings have been inconsistent across different allergic outcomes and ages of assessment: studies report protective effects of sibling presence against food allergy at 1 year^12^ and asthma at 6 years^15^ while others find no significant associations between sibling presence and food allergy at 1–2 years^14^ or multiple allergic conditions at 5 years^21^ despite documented microbiota changes. This conflicting evidence may partly reflect the difficulty in separating these interrelated pathways, heterogeneity in outcomes, and developmental timing, and/or varying sample sizes and statistical power, highlighting the need for approaches that can distinguish direct sibling effects from confounding by maternal reproductive history.

In this study, we use a large longitudinal birth cohort to investigate patterns of infant microbiome development associated with parity and sibling presence. Using triadic sampling (mother–father– infant) from over 800 families in the Finnish Health and Early Life Microbiota (HELMi) cohort^22^, we characterise compositional differences, maturation patterns, and family-level microbiota structures associated with parity/sibling presence. We examine signatures consistent with social transmission (such as within-family microbiota similarity and associations with both parental microbiota) alongside patterns that could reflect maternal reproductive history (such as timing-dependent effects in mothers). By combining >1,200 cross-sectional parental and >6,000 longitudinal infant microbiota profiles from the first two years of life with comprehensive family and health metadata, we aim to characterise the nature and extent of parity/sibling -associated microbiota differences across the family unit.

## Methods

### Cohort data

A total of 1,055 families were recruited to the Health and Early Life Microbiota (HELMi) cohort (NCT03996304) during 2016–2018 from the general population in the Helsinki metropolitan area, Finland. Only healthy, singleton babies born healthy on gestational weeks 37–42 with a body weight >2.5 kg were included in the cohort. Recruitment and data collection are described in detail in Korpela et al., 2019^22^. Briefly, faecal samples were collected from both parents at a single timepoint shortly before or after delivery and at several timepoints from infants during the first two years (at 3, 6, and 12 weeks, as well as 6, 9, 12, 18, and 24 months). Electronic questionnaires were used to collect extensive data on nutrition, growth, health, environment, lifestyle, and parental factors at one or multiple timepoints (depending on the question), including information on maternal parity and sibling household composition. Information related to delivery (mode of delivery, administration of intrapartum antibiotics, maternal group B streptococcus status, and timing of membrane rupture) was acquired from the hospital records. Additional detailed data on reproductive history and sibling characteristics (including delivery details and potential changes in living arrangements) were collected in May 2023 using an online questionnaire. Out of all 1,055 families, 346 families participated in this additional data collection. In the present study, for analyses involving infants only vaginally delivered infants (*n*=844) were included. Out of these families, 293 participated in the additional data collection. Collected data is described in detail in Jokela et al., 2023^13^.

### Ethics statement

The HELMi study was approved by The Hospital District of Helsinki and Uusimaa and conducted in accordance with the principles of the Helsinki Declaration. Participating parents signed an informed consent at enrolment and retained the right to withdraw at any point.

### DNA extraction, library preparation, and amplicon sequencing

Faecal sample processing of HELMi samples for microbiota analysis is described in detail in Jokela et al. 2023^13^. In brief, DNA was extracted from the faecal samples using a repeating bead-beating step, followed by amplicon sequencing targeting the V3–V4 region of the 16S ribosomal RNA gene. Amplicons were sequenced using Illumina MiSeq and HiSeq sequencing technology at the Functional Genomics Unit and Institute for Molecular Medicine Finland, University of Helsinki (Helsinki, Finland).

### Data processing

Data processing and subsequent analyses were conducted using R v4.1.2. Demultiplexed sequencing reads were processed through the DADA2 pipeline^23^. Taxonomy was assigned using the SILVA v.138.1 database^24^. Only ASVs assigned to kingdom Bacteria were included, and ASVs assigned to order Chloroplast or family Mitochondria were excluded. The presence of any potential contaminants using the R package *decontam*^25^ using the `prevalence` method, in which the prevalence of each sequence in biological samples is compared to that in negative controls. An ASV was considered a contaminant if it reached a probability of 0.1 in the Fisher’s exact test used in *decontam*. 15 ASV contaminants were identified and filtered from the data before subsequent analysis. The package *iNEXT* was used to generate sample type and sequencing platform -specific sample completeness and rarefaction curves, the plateau points of which were used to determine a threshold for minimum read depth for each sample type across the two different sequencing platforms. Samples with a sequencing depth under this threshold were excluded, resulting in a dataset of 1009 unique families. Taxa were then aggregated at genus level to account for variable sequencing quality and platform differences, followed by measurement of asymptotic genus-level richness and Shannon diversity using *iNEXT*^26^, which is applicable across taxonomic resolutions, including OTUs, ASVs, and higher taxa, and thus suited our genus-level data structure. The R package *phyloseq*^27^ was then used to normalise genus counts to proportional abundances.

### Analysis

All analyses were conducted in R version 4.3.1. Parity/sibling presence (highly correlated) was assessed qualitatively unless stated otherwise. To assess overall compositional differences, we performed permutational (*n*=999) multivariate analysis of variance (PERMANOVA) using genus-level Jaccard dissimilarity matrices using `adonis2` function from R package *vegan*^28^. Models were fitted separately for each timepoint, adjusting for relevant technical and biological covariates. Homogeneity of dispersion between groups was tested using `betadisper` function. To identify taxa distinguishing infants from different family structures (siblings vs no siblings), we trained Random Forest classification models for each timepoint using genus-level relative abundances (*randomForest* R package^29^). Model performance was evaluated using classification accuracy. Variable importance was assessed using mean decrease Gini index.

To test whether infants with similar family structures showed greater microbiota similarity, we calculated pairwise Jaccard dissimilarities between all infant pairs at each timepoint. Bayesian regression models (using function `brm` from R package *brms*^30^) tested whether dyadic sibling concordance (both with siblings, both without siblings, or discordant) predicted microbiota similarity, adjusting for covariates.

### Microbiota maturation

To assess whether sibling presence was associated with the maturation of the gut microbiota in early life, we predicted infant age using a Random Forest regression model (R package *randomForest*^29^) trained on genus-level relative abundances. Sequencing platform was strongly confounded with infant age (early timepoints predominantly sequenced on HiSeq, later timepoints on MiSeq). To account for this technical variation, sequencing platform was included as a binary predictor variable (MiSeq vs HiSeq) in the Random Forest model, allowing the model to learn age-related patterns while adjusting for batch effects. The model’s predictive performance was assessed using 5-fold cross-validation, where each sample was used for testing exactly once. To test parity effects on microbiota maturation, we fitted a generalised additive model (GAM) using the *mgcv* R package. Here, estimated microbiota age was used as the response variable. A smooth term for chronological age was allowed to vary by sibling presence, testing whether developmental trajectories differed between groups. The model also included other predictors, such as sequencing platform and type of feeding. Family ID was included as a random effect to account for repeated measures. Additional models tested dose-response relationships by restricting analyses to families with 1–2 older siblings and testing whether sibling number or youngest sibling age predicted microbiota age.

To identify when during development parity effects on microbiota age were most pronounced, we fitted `brm` models (*brms* package^30^) separately for each timepoint, with microbiota age as the outcome and sibling presence (yes/no) as the predictor, adjusting for covariates. To identify key taxa underlying parity-associated maturation differences, we used a ‘drop-one-taxon’ approach where each genus was dropped at a time and the age prediction model was refit^31^. Genera whose removal substantially degraded model performance were considered key contributors to age prediction. Age trajectories of these key genera were then visualised using GAMs, with smooth terms for chronological age allowed to vary by sibling presence.

### Microbiota convergence with adult samples

Microbiota similarity between infant–parent dyads was calculated using Jaccard dissimilarity at each infant timepoint (parental samples were collected around the time of the birth of the index child such that infant–parent sample pairs were not temporally matched). Bayesian regression models (using function `brm` from *brms*^30^) tested whether parity (continuous: number of previous deliveries/siblings) predicted infant–parent similarity, adjusting for covariates. Models were fitted separately for each infant timepoints and for infant–mother and infant–father dyads. To test whether convergence reflected similarity to own parents versus adults generally, we compared infant–adult dyads from the same vs different families, testing whether family membership and family composition concordance (both multi-child, both single-child, or mixed) predicted similarity).

For all `brm` models, posterior checks were conducted to confirm reliable model performances. Here, we ensured that the chains had converged, Rhat values were <1.05, bulk effective sample sizes at least 100 times the number of chains and no smaller than 10% of total posterior draws, and that the sampler took small enough steps to avoid excess (>10) divergent transitions after warm-up. Additionally, posterior predictive checks were performed to assess model fit, confirming that the simulated posterior distributions closely matched the observed data, with only minor deviations in certain regions.

### Metacommunity analyses

For each infant–mother–father triad, we calculated the total number of unique genera (family metacommunity size) and the proportion of genera unique to each family member (this analysis was repeated for each infant sample type). Differences between multi-child and single-child families were tested at each timepoint using Wilcoxon rank sum tests with FDR correction across timepoints. To test overall effects across the developmental trajectory, we fitted GAMs with the proportion of unique genera as the outcome, including smooth terms for infant age (allowing trajectories to vary by parity group) and a parametric term for parity, with family ID as a random effect (*mgcv* R package). Separate models were fitted for infant, mother, and father proportions. To test whether multi-child family metacommunities represent subsets of single-child family metacommunities, pairwise Jaccard dissimilarity between triads was decomposed into nestedness and turnover components using the *betapart* R package^32^. For each timepoint, dissimilarities were compared within vs across parity groups using Wilcoxon rank sum tests (FDR corrected across timepoints). To identify specific genera contributing to parity-associated metacommunity differences, genus-level presence frequencies were compared between parity groups at each timepoint using Fisher’s exact tests. P-values were adjusted using FDR correction across all genus–timepoint combinations.

## Results

### Sibling presence is associated with subtle compositional differences but does not create microbial homogenisation among infants

During the first weeks of life (at 3 and 6 weeks), sibling presence (highly correlated with maternal parity; assessed qualitatively) showed no significant association with genus richness (as indicated by credible intervals (CIs) overlapping zero in Bayesian regression models (brm)). Positive associations were observed at 3 months (posterior mean 0.27, CIs 0.13 to 0.42), 6 months (posterior mean 0.14, CIs 0.04 to 0.25), 18 months (posterior mean 0.09, CIs 0.02 to 0.16), and 2 years (posterior mean 0.09, CIs 0.01 to 0.16), with associations at the latter timepoints being marginal. No significant associations were observed at 9 months or 1 year (Table SX). Shannon diversity (reflecting community evenness) showed no significant associations with sibling presence at any timepoint.

To assess whether sibling presence was associated with overall shifts in community composition, we performed multivariate marginal permutational ANOVA (PERMANOVA) analyses on Jaccard distance across infant timepoints, adjusting for technical and biological covariates (covariates were determined separately for each infant timepoint based on whether they predicted microbiota variation at that timepoint). Sibling presence showed significant associations with genus-level community composition at all timepoints, though the proportion of explained variance was small (R^2^<1.0% and *p*<0.005 for all) and significantly different within-group variances were observed for several timepoints (3 weeks, 18 months, 24 months). This suggests that observed results at these timepoints may partly reflect heterogeneity in dispersion rather than true differences in community structure in infants with or without siblings.

To assess whether specific taxa could reliably distinguish between infants from different family structures, we trained Random Forest classification models for each timepoint using genus-level microbiota composition. These models achieved modest accuracy (56–71% across timepoints) in distinguishing infants from single-child vs multi-child families. Although classification performance was limited, certain taxa – particularly *Bifidobacterium, Flavonibacterium,* and *Clostridium sensu stricto 1* – consistently ranked highly in importance in early, mid-, and late timepoints, respectively, suggesting a potential temporal shift in the taxa most strongly associated with siblings.

Whilst PERMANOVA indicated subtle compositional differences between groups, this does not establish whether infants with siblings share a distinctive ‘sibling microbiota’ signature. To test whether infants from families with similar structures showed greater microbial similarity to each other than to infants from different family structures, we compared microbiota composition between infant pairs across families at each timepoint, examining whether dyads with concordant sibling background (where both had or both lacked siblings) were more similar than dyads with discordant sibling background. Across all timepoints, dyadic sibling background (yes–yes / no– no / yes–no) was not a significant predictor of microbial similarity (brm models).

To assess whether parity/sibling -associated microbiota patterns extended beyond infants, we tested for compositional differences in maternal and paternal samples. In fathers, sibling presence was significantly associated with microbiota composition (PERMANOVA; R^2^=0.004, F=1.793 *p*=0.011; beta dispersion F=2.274, *p*=0.139; *n*=552). In mothers, however, sibling presence effects depended on sampling time relative to birth (no such effect observed in fathers). When pre- and post-birth maternal samples were analysed together, no significant association was detected (R^2^=0.002, F=1.417, *p*=0.071; beta dispersion F=1.827, *p*=0.196; *n*=701), though significant differences in within-group dispersion were observed between pre- and post-birth samples (beta dispersion F=7.848, *p*=0.004). Stratifying by sampling time revealed that sibling presence effects were only detectable in samples collected after giving birth (R^2^=0.008, F=1.780 *p*=0.008; beta dispersion F=0.783, *p*=0.381; *n*=488) but not before birth (R^2^=0.003, F=1.453, *p*=0.045; beta dispersion F=9.042, *p*=0.005; *n*=213).

### Parity accelerates microbiota maturation

To assess whether parity was associated with the rate of gut microbiota maturation in early life, we predicted infant age using a Random Forest regression model trained on genus-level relative abundances across eight timepoints from birth to 2 years. The predicted ‘microbiota age’ was strongly correlated with chronological age (cross-validated R^2^=0.853, RMSE=6.092 months; Fig. 1A). Sequencing platform was included as a predictor to account for strong confounding with infant age (Table S1) and ranked 19^th^ out of the 77 features in variable importance, with the top predictors being genera associated with microbiota maturation (*Anaerostipes, Faecalibacterium, Blautia*; Table S1).

**Figure 1.**
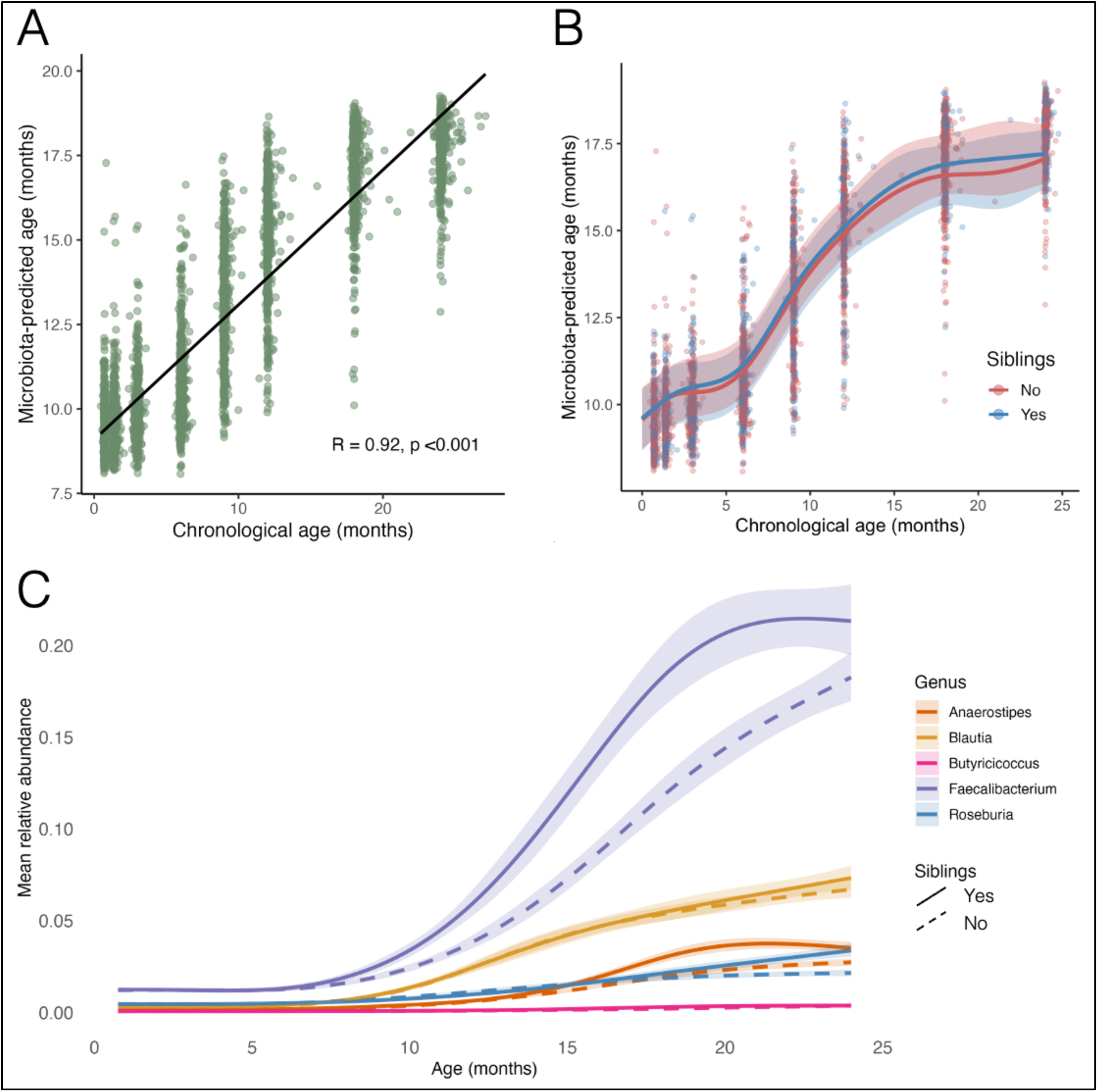
(**A**) Relationship between chronological age and microbiota-predicted age in infants aged 3 weeks to 2 years, modelled using a Random Forest regression model trained on genus-level relative abundance data. Cross-validated R^2^=0.853, RMSE=6.092 months. Model performance was assessed using 5-fold cross-validation, where each sample was used for testing exactly once. Model stability was further supported by Out-of-Bag (OOB) performance metrics (average OOB R²=0.879 ± 0.002, OOB RMSE=8.06 ± 0.09 months). (**B**) Generalised additive model predictions showing microbiota age trajectories for infants with (blue) and without (red) older siblings. Lines show fitted values with 95% confidence intervals; points show observed data. (**C**) Generalised additive model visualising top 5 genera with highest importance scores in microbiota age prediction in infants from multi-child (solid lines) or single-child (dashed lines) families.

We hypothesised that infants with siblings would experience acceleration of microbiota maturation (where microbiota age is higher than expected from chronological age) due to early exposure to microbiota from older siblings, effectively acquiring microbes representative of later stages of microbiota development. In support of this, a generalised additive model (GAM) showed distinct microbiota age trajectories for infants with and without siblings (*p*<0.001 for both smooth terms; Fig. 1B), indicating a modest acceleration of maturation in infants from multi-child families (estimate=0.140, *p*=0.002; *n*=4,305 from 808 infants; 1–8 samples per infant; Table S2). To test whether this effect was modified by the number of older siblings or the age of the youngest sibling, we fitted additional models. When restricting to infants from families with one or two other children (*n*=1,657 samples from 304 infants; 249 infants with one sibling, 55 infants with two siblings), the number of siblings did not significantly predict microbiota age (estimate=0.107, *p*=0.219). Further, in a subset with data on sibling age, the age of the youngest older sibling did not significantly predict microbiota age (*n*=851 samples from 147 infants with 1–2 siblings; estimate=0.003, *p*=0.123).

To examine when parity differences in microbiota maturation were most pronounced, we ran brm models separately for each time point and found that sibling presence (yes/no) was significantly associated with elevated microbiota age at 6 (posterior mean 0.03, CIs 0.01 to 0.05), 9 (posterior mean: 0.05, CIs 0.01 to 0.09) and 18 months (posterior mean: 0.02, CIs 0.01 to 0.03; Fig. 1B), although the effects were small and only marginally significant. Other available measures of social transmission (number of caregivers, day care attendance) did not predict microbiota age (these predictors were only included in models with infants of 1–2 years). Early-life feeding was significantly associated with microbiota age at several timepoints such that lack of breastfeeding (be that use of formula as an alternative or absence of either milk in diet) accelerated microbiota maturation, with effects consistently larger than those of presence of siblings.

To better understand which microbial taxa underlie microbiota age predictions, we investigated the age trajectories of the five genera whose removal most increased prediction error. Several taxa revealed distinct age trajectories between infants with and without siblings (Fig. 1C). Notably, *Faecalibacterium –* the second most important genus in predicting microbiota age – exhibited an earlier and more rapid increase in relative abundance among infants with siblings compared to those without. While the *Faecalibacterium* trajectory appeared to plateau at around 20% relative abundance in infants from multi-child families, the single-child group showed a continued upward trend without a clear plateau within the observed sampling window (Fig. 2C).

**Figure 2.**
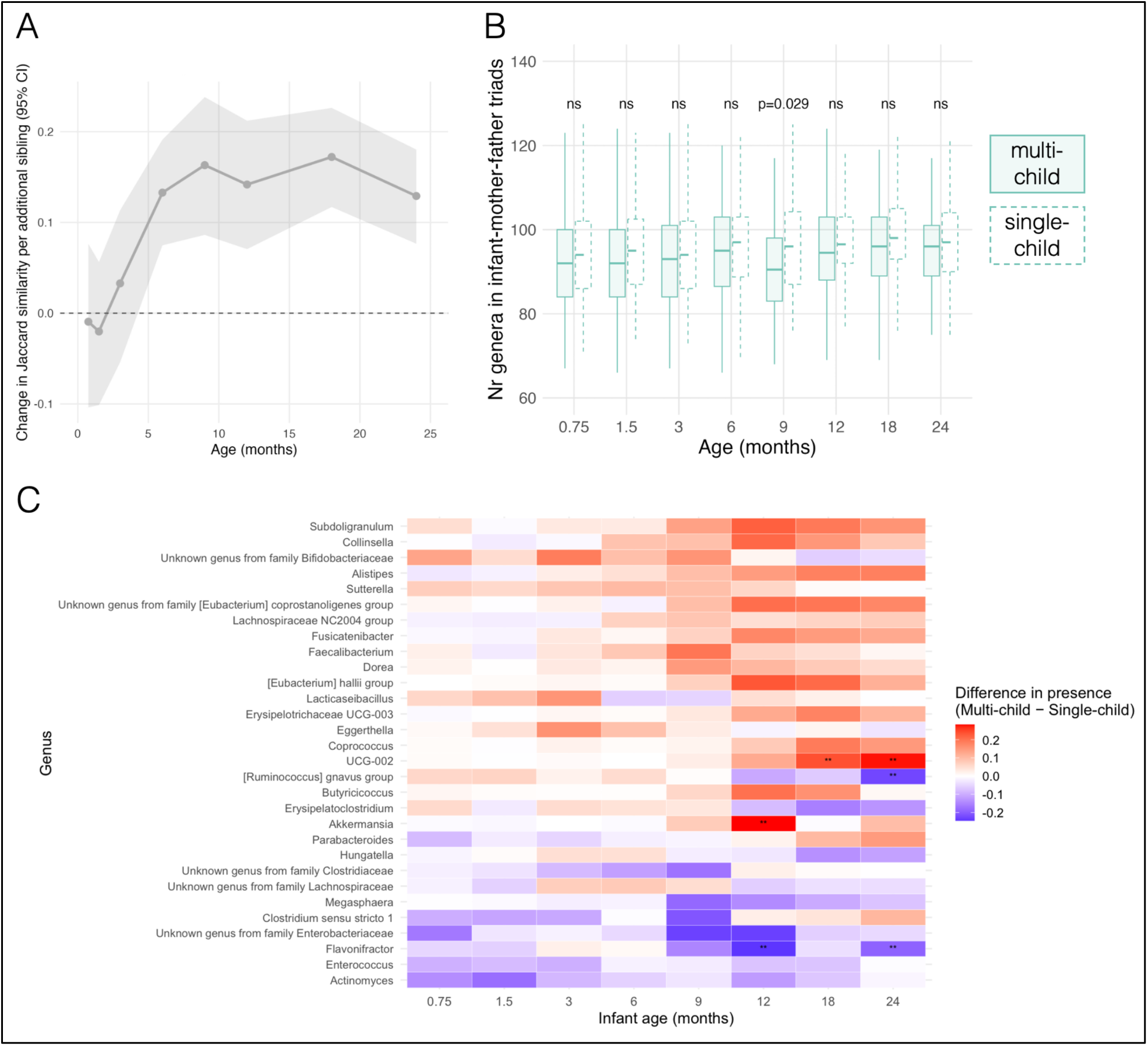
(**A**) Parity effects on infant-mother similarity across infant ages. Points show the estimated effect of parity (posterior means with 95% credible intervals (CI)) from separate Bayesian regression models fitter at each timepoint. The y-axis represents the change in similarity per additional previous pregnancy (i.e., number of older siblings). **(B)** Number of unique genera in infant–mother–father triads from multi-child (solid boxes) vs single-child (dashed boxes) families. Number of triads varied from 224 to 319 across the 8 timepoints. Statistical differences between multi-child and single-child families were tested with Wilcoxon rank sum tests (FDR corrected). Ns, *p*>0.05. **(C)** Genus-level differences in presence frequency between multi-child and single-child families across infant ages. For each infant age group, presence frequency of each genus was compared between triads from multi-child and single-child families using Fisher’s exact tests. Genera shown represent the 30 with the highest average absolute frequency difference across timepoints. Tile colour indicates the magnitude and direction of the difference in presence frequency (multi-child – single-child). After correction for multiple comparisons across all genus-timepoint combinations (*n*=3,504 tests), only 6 comparisons remained significant (FDR *p*<0.05). **, p<0.01.

### Microbiota of infants with older siblings converge more with adults

Across infant–parent dyads (infant–mother or infant–father), higher parity/number of siblings (modelled as a continuous predictor) was significantly associated with greater microbiota similarity from 3 months of infant age onwards (Fig. 2A, Fig. S1; parental samples were collected shortly before or after index child’s birth, such that samples in dyads were not temporally matched). At 3 months of age, having 1–2 in contrast to >2 caregivers increased similarity in infant–parent dyads (brm models; infant–father: posterior mean 0.54, CIs 0.21 to 0.92; infant–mother: posterior mean 0.28, CIs 0.02 to 0.56; families with 1–2 caregivers; families with >2 caregivers) as expected when fewer competing microbial sources are present. In infant– father dyads, this smaller number of caregivers increased microbiota similarity also at 3 and 6 weeks of age (3 weeks: posterior mean 0.31, CIs 0.03 to 0.62; 6 weeks: posterior mean: 0.38, CIs 0.05 to 0.74). Day care attendance nor day care group size (only assessed at 1.5 and 2 years of age when sufficient variation was available) predicted infant–parent similarity.

To assess whether this convergence reflects greater similarity to one’s own parent vs adult microbiotas more generally, we compared infant–adult dyads from the same and different families. Family composition similarity (both multi-child, both single-child, or mixed) significantly predicted microbiota similarity at several timepoints across dyads (brm models), whereas family membership had a detectable effect only at 18 months, and this effect was weaker than that of parity. This suggests that parity promotes convergence with adult-like microbial profiles broadly, rather than uniquely increasingly resemblance to the specific parent. In contrast to infant–parent dyads, parity did not increase microbial similarity in mother–father dyads (posterior mean: 0.05, CIs −0.02 to 0.12).

We next investigated whether the accelerated microbiota maturation in infants from multi-child families was associated with acquisition of a broader range of taxa, which could manifest as larger metacommunities within infant–mother–father triads. Contrary to this, multi-child families consistently showed smaller metacommunities than single-child families across all timepoints, although this difference was statistically significant only at 9 months of infant age (Wilcoxon rank sum test, FDR-corrected *p*=0.029; *p*>0.05 at remaining timepoints; Fig. 2B). The proportion of genera unique to infants (relative to the family-level metacommunity) was marginally lower in multi-child families across the developmental trajectory (GAM parametric coefficient=-0.004, *p*=0.050). A similar pattern was observed for the proportion of genera unique to mothers (coefficient=-0.008, *p*=0.035), whereas father showed no significant difference (coefficient=0.002, *p*=0.634). These proportions were calculated at each infant timepoint relative to the metacommunity of that infant–mother–father triad.

To investigate whether metacommunities in multi-child families represent subsets of the larger metacommunities observed in single-child families, we decomposed pairwise Jaccard dissimilarity among infant–mother–father triads into components of nestedness (asymmetric subset relationships) and turnover (taxa replacement). For each infant age group, we compared dissimilarities within the same parity group (‘within’) and across different groups (‘between’). Turnover was significantly higher in cross-parity comparisons than within-parity comparisons at 7 of 8 timepoints (Wilcoxon rank sum test, FDR-corrected *p*<0.05; exception at 6 months, *p*=0.876; Fig. S2A), indicating that triads from different sibling contexts harbour taxonomically distinct sets of genera throughout development. Turnover declined with infant age in both comparison types, from approximately 65% at 3 weeks to 50% at 24 months (Fig. S2A), reflecting developmental convergence toward more similar genus compositions across families. However, the taxonomic distinctiveness between parity groups persisted despite this overall convergence. In contrast, nestedness was generally low (median<0.1) and did not differ significantly between groups, except at 9 months, where a small but statistically significant difference was detected (FDR-corrected *p*=0.033; Fig. S2B). The higher turnover between groups indicates that multi-child and single-child family metacommunities contain different sets of genera rather than one being a subset of the other.

To explore which genera showed the largest differences in occurrence between parity groups, we examined genus-level presence across infant–mother–father triads, comparing frequency between parity groups at each timepoint. After correction for multiple comparisons across all genus-timepoint combinations, only 6 of 3,504 tests remained significant (FDR *p*<0.05). We visualised the top 30 genera showing the largest average differences in presence frequency across ages (Fig. 2C). *Akkermansia* was significantly more frequent in multi-child family triads at 12 months (50% vs 22%, FDR-corrected *p*=0.008), while *Flavonifractor* was significantly less frequent in multi-child families at both 12 months (66% vs 91%, *p*=0.007) and 24 months (73% vs 93%, *p*=0.007).

## Discussion

In this study, we characterised the effects of parity and sibling presence on infant gut microbiota development using triadic sampling from over 800 families. While parity/sibling presence showed significant associations with microbiota composition across infants, mothers, and fathers, the effects were modest. Infants from multi-child families exhibited accelerated microbiota maturation and enhanced convergence with adult microbiota, yet infants from similar family structures did not show increased microbial similarity to one another. Multi-child families harboured smaller but compositionally distinct metacommunities, with differences driven by taxonomic turnover rather than loss of diversity. Together, these findings suggest that family structure influences infant microbiota throughout multiple pathways that may involve both maternal reproductive biology and household-level factors.

Direct biological effects of parity are expected in mothers, yet parity/sibling-related patterns were detected in both mothers and fathers. This suggests that at least some parity/sibling-associated microbiota patterns reflect household-level factors (such as changes in diet^16^ or hygiene practices^20^, or social transmission of microbes^10^) that affect the entire family rather than maternal biology alone. The timing-dependent pattern in mothers (effects only detectable in post-birth sampled mothers) could reflect pregnancy-related changes in the maternal gut microbiota^33^, though longitudinal sampling would be needed to confirm this.

The modest compositional associations we observed are difficult to reconcile with a straightforward sibling transmission model. While sibling presence explained small but significant variance in community composition at all timepoints, suggesting subtle directional shifts rather than distinctive community structures, these effects did not translate to increased microbial similarity amongst infants from similar sibling backgrounds. Existing literature has interpreted associations between sibling presence and infant microbiota composition as evidence of direct sibling–to–infant microbial transmission^14^. However, without microbiota profiles from the siblings themselves, such conclusions remain speculative. Observing that infants with siblings have different microbiota than only children does not establish that siblings are the source of those microbes – it is equally consistent with confounding by maternal reproductive history, differences in household practices, or other family-level factors associated with having multiple children. If siblings were a major source of shared microbes, we might expect infants with siblings to resemble each other more than they resemble only children. The lack of such homogenisation in our data suggests either that sibling transmission occurs but involves highly individualised taxa (reflecting heterogeneity in siblings’ own microbiota across different ages and developmental stages), or that parity/sibling -associated factors beyond direct sibling contact drive these compositional differences.

We observed modest but significant acceleration of microbiota maturation in infants with siblings, aligning with previous studies^4,12^. Infants from multi-child families showed elevated microbiota age, particularly at 6, 9 and 18 months, with *Faecalibacterium* – a key predictor of microbiota age – exhibiting earlier and more rapid increases. Consistent with accelerated maturation, these infants also showed increased genus richness at several timepoints. Shannon diversity showed minimal associations, suggesting compositional shifts involve changes in presence/absence of taxa rather than substantial restructuring of community evenness. Sibling transmission appears a plausible mediator of these effects, as older siblings would provide exposure to both more developmentally advanced microbiota (potentially accelerating maturation) and a broader range of taxa (increasing richness). However, when we examined whether this effect scaled with the number of siblings (comparing infants with one vs two siblings) or with the age of the youngest older sibling, neither factor significantly predicted microbiota age. While these analyses were limited by sample size and may have lacked statistical power, the lack of dose-response or age-dependent effects raises questions about direct sibling transmission. Several alternative mechanisms could contribute to this pattern. Parity-related changes in maternal gut microbiota^18,19^ could facilitate earlier establishment of taxa like *Faecalibacterium* through vertical transmission at birth or ongoing post-birth maternal transmission. Thus, while maturation acceleration and associated richness increases are consistently associated with sibling presence, the mechanisms remain ambiguous.

Consistent with accelerated maturation, infants from multi-child families also showed increased convergence with adult microbiota profiles. Parity/sibling presence was associated with increased infant–parent similarity from 3 months onwards, yet this convergence occurred with adult microbiota broadly rather than being specific to the infant’s own parents. Importantly, mother–father similarity did not increase with parity/sibling presence, suggesting that the observed pattern reflects changes in infant microbiota rather than household-level effects. Adult microbiota is considerably more stable than infant microbiota due to mature immune selection and established gut physiology, making it more resistant to colonisation by exogenous microbes^34^. The non-specific nature of this convergence (where infants become more similar to adults generally rather than their own parents specifically) reinforces that these infants are maturing more rapidly towards adult-like profiles. This pattern could arise if sibling presence increases infant exposure to adult microbiota through expanded social networks (visitors, other parents at activities), facilitating transmission of adult-type taxa that the infant’s developing gut can accommodate. Alternatively, parity-related changes in maternal physiology, immunity, or microbiota composition could alter the infant gut environment in ways that favour establishment and persistence of adult-type taxa, either through modified vertical transmission or through programming of selective pressures in the infant gut. These mechanisms are not mutually exclusive, and both could contribute to the observed acceleration of maturation and convergence with adult profiles.

The family-level metacommunity analysis provides additional insight into how family structure shapes microbial communities. Multi-child families harboured smaller metacommunities than single-child families, with differences driven by taxonomic turnover rather than nestedness, meaning these families support compositionally different assemblages rather than simply reduced subsets of single-child family taxa. This reduction in metacommunity size was driven by infants contributing fewer unique genera relative to their family’s metacommunity, while parental contributions remained similar across family types. This pattern indicates that infants in multi-child families harbour microbiota that overlap more extensively with their parents’ communities, consistent with the accelerated convergence towards adult-like profiles described above. Whether this increased overlap reflects more efficient transmission of parental taxa or these infants developing selective pressures that favour adult-type taxa remains unclear. However, the observation that specific (even if only a few) genera show substantially different presence frequencies between family types (e.g., *Akkermansia* more common in multi-child families, *Flavonifractor* in single-child families) suggests that family structure creates distinct selective pressures favouring different taxa. If sibling presence primarily increased microbial diversity through transmission, we might expect multi-child families to harbour additional taxa on top of those found in single-child families, rather than the compositional turnover and reduced metacommunity size we observed.

Our findings have implications for interpreting epidemiological associations between sibling presence and health outcomes. Previous studies have attributed protective effects against allergies to increased microbial exposure from siblings^12,15^, yet our results suggest that such effects, if real, may operate through pathways beyond direct sibling transmission. If maternal reproductive history independently shapes infant microbiota – through alterations in maternal microbial communities, immune factors in breastmilk, or parenting practices – then the causal pathways to health outcomes may differ substantially from those assumed. This could explain inconsistent findings regarding sibling presence and allergic outcomes^14,21^, as different studies may capture different aspects of this complex relationship depending on their ability to account for maternal factors.

Several limitations warrant consideration. While our triadic design provides leverage for distinguishing household-level from maternal-specific effects, parity and sibling presence remain correlated, limiting our ability to fully separate their influences. As we lack microbiota profiles from siblings, we cannot directly assess sibling-to-infant microbial transmission. Our analyses used 16S rRNA amplicon sequencing data, which provides genus-level resolution but cannot detect strain-level transmission patterns. Our study is observational, precluding definitive causal inference about the mechanisms generating observed patterns. While the data comprises >800 family triads, sample sizes for detailed reproductive history analyses were modest, limiting statistical power for detecting associations with specific pregnancy outcomes such as known miscarriage. Our cohort is relatively homogenous in socioeconomic status and findings may not generalise to populations with different environmental and cultural contexts.

In conclusion, while sibling presence is associated with infant microbiota differences, our findings suggest these effects likely reflect multiple interacting factors rather than primarily direct sibling transmission. The lack of increased similarity amongst from similar sibling backgrounds, the non-specific convergence with adult microbiota, and the compositionally distinct metacommunities in different family contexts point towards more complex mechanisms that may maternal reproductive history, household practices, and family-level dynamics.

## Author information

### Contributions

EH and AS conceptualised the presented study. EH and AJP processed the sequencing data. EH and AJP curated the metadata. EH analysed the data. EH wrote the manuscript. AS, KLK and WMdV conceived the HELMi cohort, obtained funding, and provided data access. All authors contributed to the final version of the manuscript.

## Data availability

The 16S rRNA amplicon sequencing data from the HELMi birth cohort is publicly available at https://www.ebi.ac.uk/ena/browser/view/PRJEB55243. Metadata, aerotolerance data, taxonomy table, ASV relative abundance table as well as R code used in this study will be made publicly available on GitHub upon publication.

## Supplementary Material

**Supplementary Figure 1.**
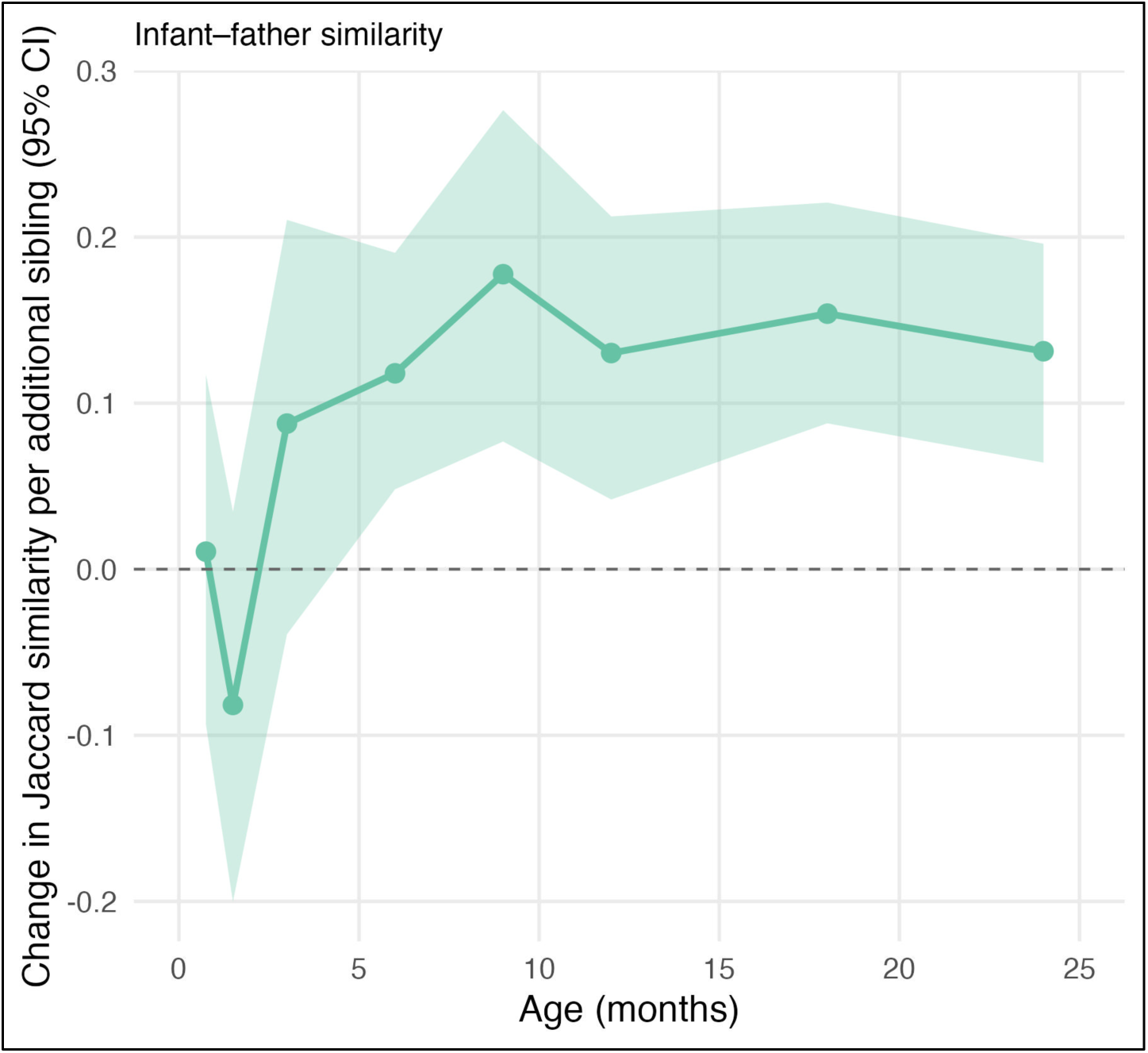
Parity effects on infant-father similarity across infant ages. Points show the estimated effect of parity (posterior means with 95% credible intervals) from separate Bayesian regression models fitter at each timepoint. The y-axis represents the change in similarity per additional previous pregnancy (i.e., number of older siblings).

**Supplementary Figure 2.**
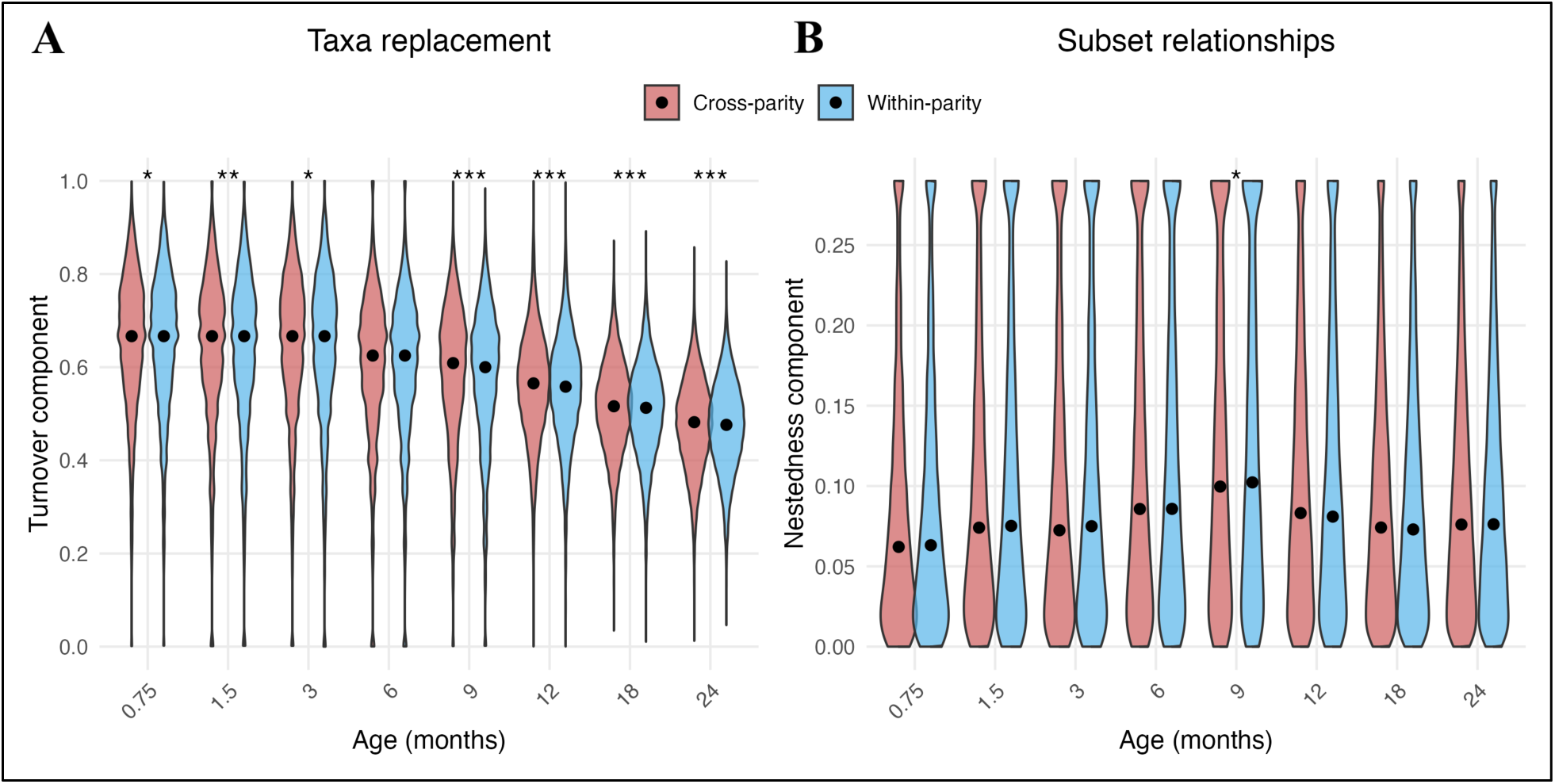
Decomposition of Jaccard dissimilarity into turnover and nestedness components for infant–mother–father triads across infant development. Pairwise dissimilarities between triads were calculated at each infant timepoint and partitioned into (**A**) turnover (taxa replacement) and (**B**) nestedness (asymmetric subset relationships) components. Comparisons are categorised as cross-parity (red; one triad from a multi-child family and one from a single-child family) or within-parity (blue; both triads from the same parity group). Violin plots show the distribution of values; black dots indicate medians. Asterisks denote significant differences between comparisons (Wilcoxon rank sum tests; *<0.05, **<0.01; ***<0.001 FDR).

**Supplementary Table 1.**
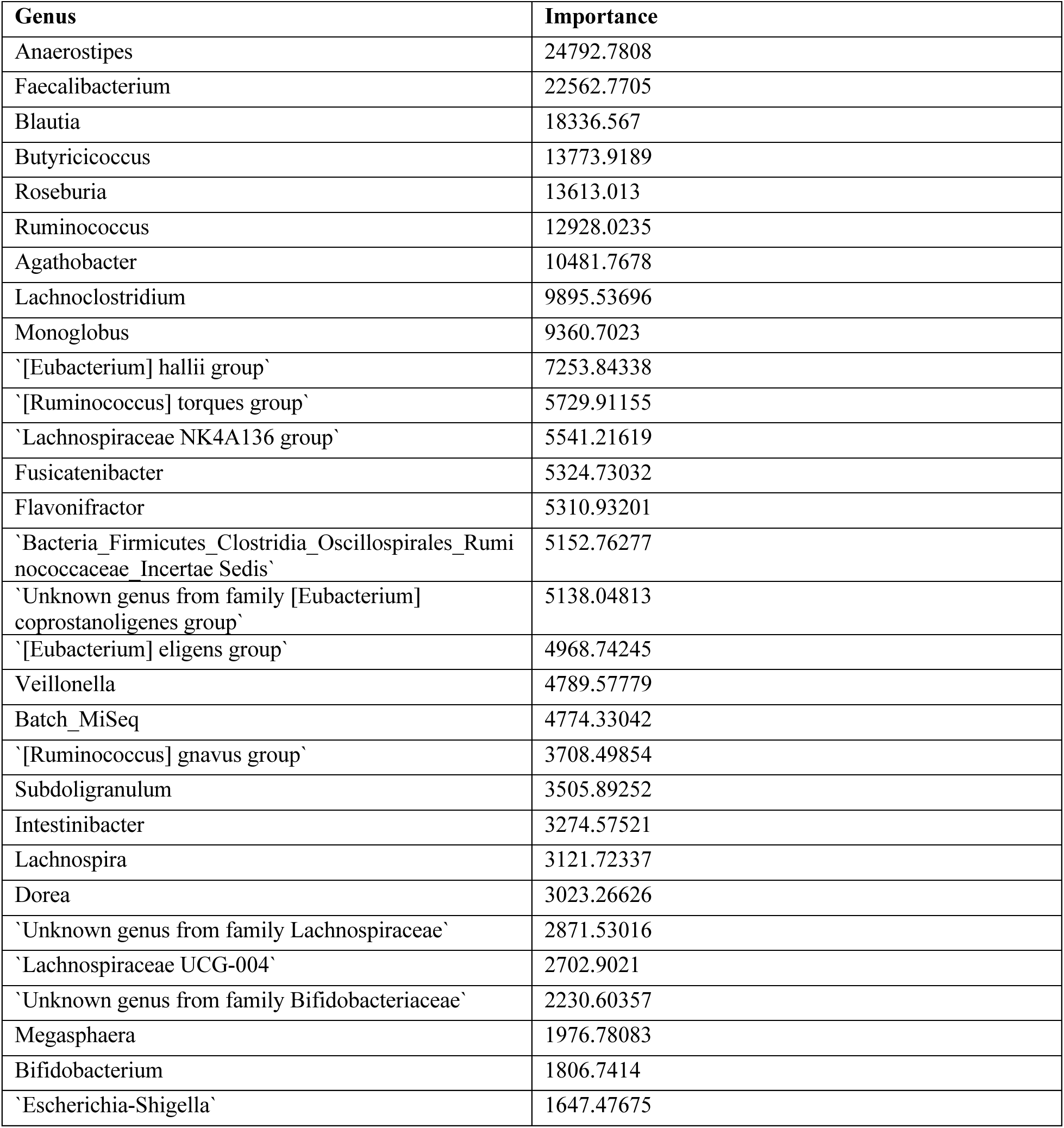
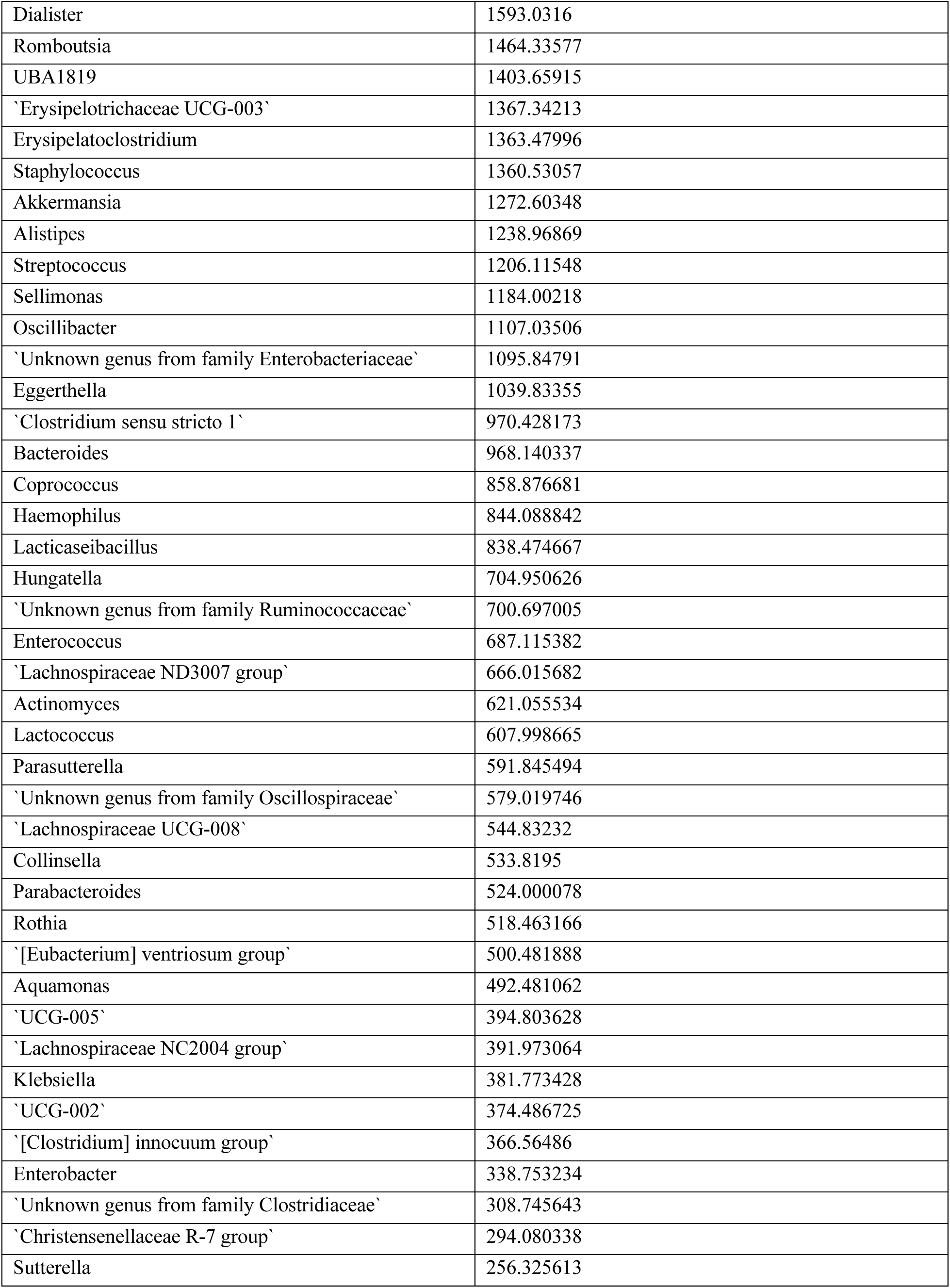
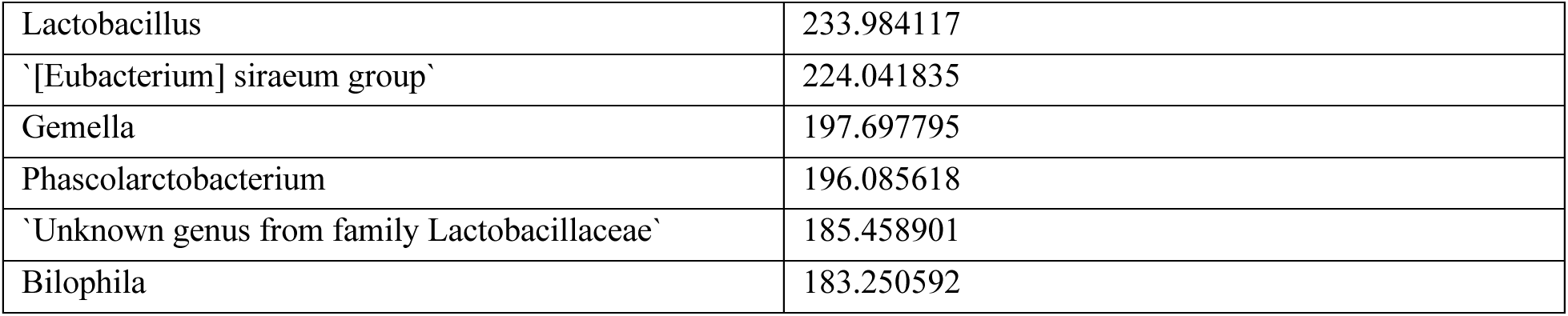
Importance scores from Random Forest regression model trained on genus-level relative abundances across predicting microbiota age. Sequencing platform was included as a binary predictor variable (MiSeq or HiSeq) as it was strongly confounded with infant age. The model’s predictive performance was assessed using 5-fold cross-validation, where each sample was used for testing exactly once (cross-validated R^2^=0.853, RMSE=6.092 months). Out-of-bag (OOB) performance metrics across folds: mean OOB R^2^=0.879 ± 0.002; mean OOB MSE=8.065 ± 0.085.

**Supplementary Table 2.** Generalised additive model predicting microbiota age from chronological age and covariates. Parametric coefficients represent linear effects on predicted microbiota age (in months). Smooth terms represent non-linear age trajectories, with estimated degrees of freedom (EDF) indicating curve complexity. The model included random intercepts for Family ID to account for repeated measures (n=4,305 samples from 808 families; 1–8 samples per infants). Reference categories: no previous deliveries (siblings), both breastmilk and formula, male infant, HiSeq sequencing platform, brown stool, no intrapartum antibiotics. Adjusted R^2^=0.918.

Parametric coefficients
| Term | Estimate | SE | t-value | p-value |
| --- | --- | --- | --- | --- |
| Intercept | 12.772 | 0.435 | 29.37 | <0.001 |
| Previous deliveries (Yes) | 0.140 | 0.045 | 3.14 | 0.002 |
| Milk: Breastmilk | -0.459 | 0.055 | -8.29 | <0.001 |
| Milk: Formula | 0.403 | 0.092 | 4.36 | <0.001 |
| Milk: None | -0.039 | 0.083 | -0.47 | 0.639 |
| Infant sex (Girl) | 0.008 | 0.043 | 0.19 | 0.849 |
| Sequencing platform (MiSeq) | 0.923 | 0.047 | 19.80 | <0.001 |
| Stool color: Clay gray | 0.229 | 0.523 | 0.44 | 0.662 |
| Stool color: Dark brown | -0.011 | 0.433 | -0.03 | 0.980 |
| Stool color: Green | -0.368 | 0.441 | -0.83 | 0.404 |
| Stool color: Light brown | -0.063 | 0.432 | -0.15 | 0.885 |
| Stool color: Yellow | -0.280 | 0.435 | -0.64 | 0.520 |
| Received intrapartum antibiotics | 0.165 | 0.051 | 3.21 | 0.001 |

Smooth terms
| Term | EDF | Ref. DF | F-value | p-value |
| --- | --- | --- | --- | --- |
| Age trajectory: No previous deliveries | 6.97 | 8 | 864.57 | <0.001 |
| Age trajectory: Previous deliveries | 6.63 | 7 | 819.36 | <0.001 |
| Family (random effect) | 377.68 | 804 | 0.93 | <0.001 |

